# Finger recruitment patterns during mirror movements suggest two systems for hand recovery after stroke

**DOI:** 10.1101/129510

**Authors:** Naveed Ejaz, Jing Xu, Meret Branscheidt, Benjamin Hertler, Heidi Schambra, Mario Widmer, Andreia V. Faria, Michelle Harran, Juan C. Cortes, Nathan Kim, Tomoko Kitago, Pablo A. Celnik, Andreas Luft, John W. Krakauer, Jörn Diedrichsen

**Author notes:** Author contributions: These authors contributed equally to this work (Naveed Ejaz, Jing Xu). Data: Data taken from (Xu *et al.*, 2016). Study design and analysis: NE, JX, JD, JWK. Manuscript: NE, JX, JD, JWK. Correspondence: Naveed Ejaz @. Open Science: Behavioural dataset available at: https://github.com/nejaz1/mirroring2017.

## Abstract

Accumulating behavioural and neurophysiological evidence suggests that upper-limb control relies on contributions from both cortical and subcortical motor circuits, with cortical inputs providing fine-finger function and subcortical inputs providing the ability for gross movements, respectively. During recovery of function after stroke, the relative contributions from these pathways may shift. Here we propose that mirror movements that appear after stroke provide a non-invasive assay through which relative contributions from cortical and subcortical pathways towards hand recovery can be studied. We hypothesized that mirror movements, like hand function, are generated by summed contributions from cortical and subcortical pathways, and suggest that subcortical contributions should be characterized by a broad recruitment of fingers, while cortical contributions primarily recruit the homologous finger in the passive hand. In a longitudinal stroke recovery study (Xu *et al*., 2016), we quantified mirror movements and paretic hand function in 53 stroke patients in the year following unilateral stroke. Mirror movements in the non-paretic hand were exaggerated early after damage (week 2), with paretic finger presses broadly recruiting multiple fingers in the non-paretic hand. On average, however, mirroring in homologous fingers was 1.76 times larger than in non-homologous fingers. Over the year, mirroring in the non-paretic hand progressively normalized with a time-course that mimicked that for the fine-finger deficits in the paretic hand. In comparison, during non-paretic finger presses, the homologous component of mirroring in the paretic hand was reduced early after stroke (week 2) but progressively normalized. Altogether, we conclude that the pattern of mirror movements across homologous and non-homologous fingers reflect the summed contributions of both cortical and subcortical systems, and we discuss the implications of our results towards hand recovery after stroke.

## Introduction

Accumulating behavioural evidence suggests that upper-limb function relies on inputs from both cortical and subcortical motor circuits. While cortical contributions towards upper-limb function are well-established (Brinkman and Kuypers, 1973; Soteropoulos *et al*., 2011), subcortical contributions have been proposed to explain why voluntary movements in response to startling acoustic cues have reaction times which are much shorter than the known conduction delays from the cortex to the upper-limb (Carlsen *et al*., 2009; Dean and Baker, 2016; Honeycutt *et al*., 2013). Furthermore, different aspects of upper-limb function (i.e. strength and fine-control) dissociate after stroke (Lan *et al*., 2017; Sukal *et al*., 2007; Xu *et al*., 2016), suggesting that these two components reflect contributions from (at least) two separate systems, originating in cortical and subcortical areas respectively (Xu *et al*., 2016).

Neurophysiological studies in primates provide additional evidence for cortical and subcortical contributions towards hand function, further suggesting that inputs from each area contribute towards different aspects of hand function. The most prominent inputs to the hand come through the corticospinal tract (Porter and Lemon, 1993; Soteropoulos *et al*., 2011), which connects motor circuits in the contralateral hemisphere to the spinal cord and provides the ability to perform fine-finger function e.g. precision grip (Lawrence and Kuypers, 1968a; Rathelot and Strick, 2009; Tower, 1940). Additional input to the hand comes from phylogenetically-older, rubrospinal and reticulospinal pathways originating in the brainstem. In contrast to the corticospinal tract, these subcortical pathways are mainly involved in gross movements (e.g. whole-hand grasping) and offer only a limited ability for fractionated finger control (Lawrence and Kuypers, 1968b; Riddle *et al*., 2009; Soteropoulos *et al*., 2012). Since the rubrospinal pathway is largely absent in man (Nathan and Smith, 1955; 1982), the reticulo- and corticospinal pathways have been proposed to mediate gross and fine-control aspects of hand function respectively (Sukal *et al*., 2007; Xu *et al*., 2016).

Together, these cortical and subcortical pathways potentially provide a certain degree of flexibility in hand function, with one partially able to compensate for damage to the other. Indeed, changes in the relative contributions of cortical and subcortical pathways in primates is one proposed mechanism through which the hand regains function following stroke (Herbert *et al*., 2015; Zaaimi *et al*., 2012). The extent to which changes in pathway contributions are responsible for hand recovery in man is unknown, primarily because invasive investigations like those in primates are not possible.

In this study, we posit that mirror movements provide a non-invasive assay through which changes in the relative contributions from cortical and subcortical systems after stroke can be studied. In health, mirror movements are unintended movements that appear in the passive hand when the active hand voluntarily moves (review, Cincotta and Ziemann, 2008). Surprisingly little is known about the nature of mirroring after stroke, except that in chronic patients they are exaggerated in the non-paretic hand (Cernacek, 1961; Y. Kim *et al*., 2015; Y. H. Kim *et al*., 2003; Nelles *et al*., 1998), but slightly reduced in the paretic hand (Nelles *et al*., 1998). While mirroring has typically been attributed to abnormally large activities in cortical sensorimotor areas (Cincotta and Ziemann, 2008; Cramer *et al*., 1997; Y. H. Kim *et al*., 2003; Ward *et al*., 2003; Wittenberg *et al*., 2000), subcortical pathways are also plausible candidates. For instance, individual reticulospinal axons project bilaterally onto the contra- and ipsilateral sections of the spinal cord (Sakai *et al*., 2009), and activate upper-limb muscles on either side of the body (Hirschauer and Buford, 2015), potentially causing mirroring.

We hypothesized that mirror movements, like hand function, might be caused by summed contributions from cortical and subcortical pathways. Furthermore, we suggest that relative contributions from these pathways can be disentangled by studying the exact patterns of finger recruitment during mirroring. Subcortical contributions to mirroring should result in a broad recruitment of fingers in the passive hand, reflecting the pathway’s limited ability to provide fractionated finger control (Lawrence and Kuypers, 1968b; Soteropoulos *et al*., 2012). In contrast, we have observed that finger presses result in activation patterns in cortical sensorimotor areas that are highly similar regardless of whether the contralateral, or the homologous finger in the ipsilateral hand was used (Diedrichsen:2013hb, also see Liu *et al*., 2010; Scherer *et al*., 2009). Therefore, cortical contributions towards mirroring should primarily recruit the homologous finger in the passive hand, reflecting the specialized role of neocortical motor areas in providing fine-finger control (Brinkman and Kuypers, 1973; Soteropoulos *et al*., 2011).

Therefore, in 53 stroke patients, we characterized the year-long changes in mirror movements after damage. After stroke, individuated finger presses with the paretic hand resulted in a broad recruitment of fingers in the non-paretic hand. On average, however, mirroring in homologous fingers was larger than in non-homologous fingers. In comparison, the homologous component of mirroring in the paretic hand was reduced early after stroke but subsequently normalized. Altogether, we conclude that mirror movements reflects contributions from (at least) two separate systems, and discuss the implications of these results on cortical and subcortical contributions towards hand recovery after stroke.

## Materials and Methods

### Participants

53 patients with hemiparesis (20 female; age=57.4, SD=14.9 years) were recruited within the first week after stroke. The recovery of paretic hand function is reported in Xu et al. (2016), but clinical measures of impairment at the time of recruitment are summarized in Supplementary Figure 1. Patients were included if they had a first-time unilateral ischemic stroke within the previous 2 weeks and reported unilateral weakness of the upper extremity (Medical Research Council muscle weakness scale<5). They were excluded if age<21 years, their initial upper-limb impairment was too mild (Fugl-Meyer>63/66), or if they had cognitive deficits that could impair task comprehension and performance. Excluding aphasic patients led to a bias of right-hemispheric infarcts (36 right), in turn leading to a disproportionately higher ratio of left-handed patients (42 right-hand; according to Oldfield (1971)). A comprehensive list of inclusion/exclusion criteria is available at Xu et al. (2016).

**Figure 1.**
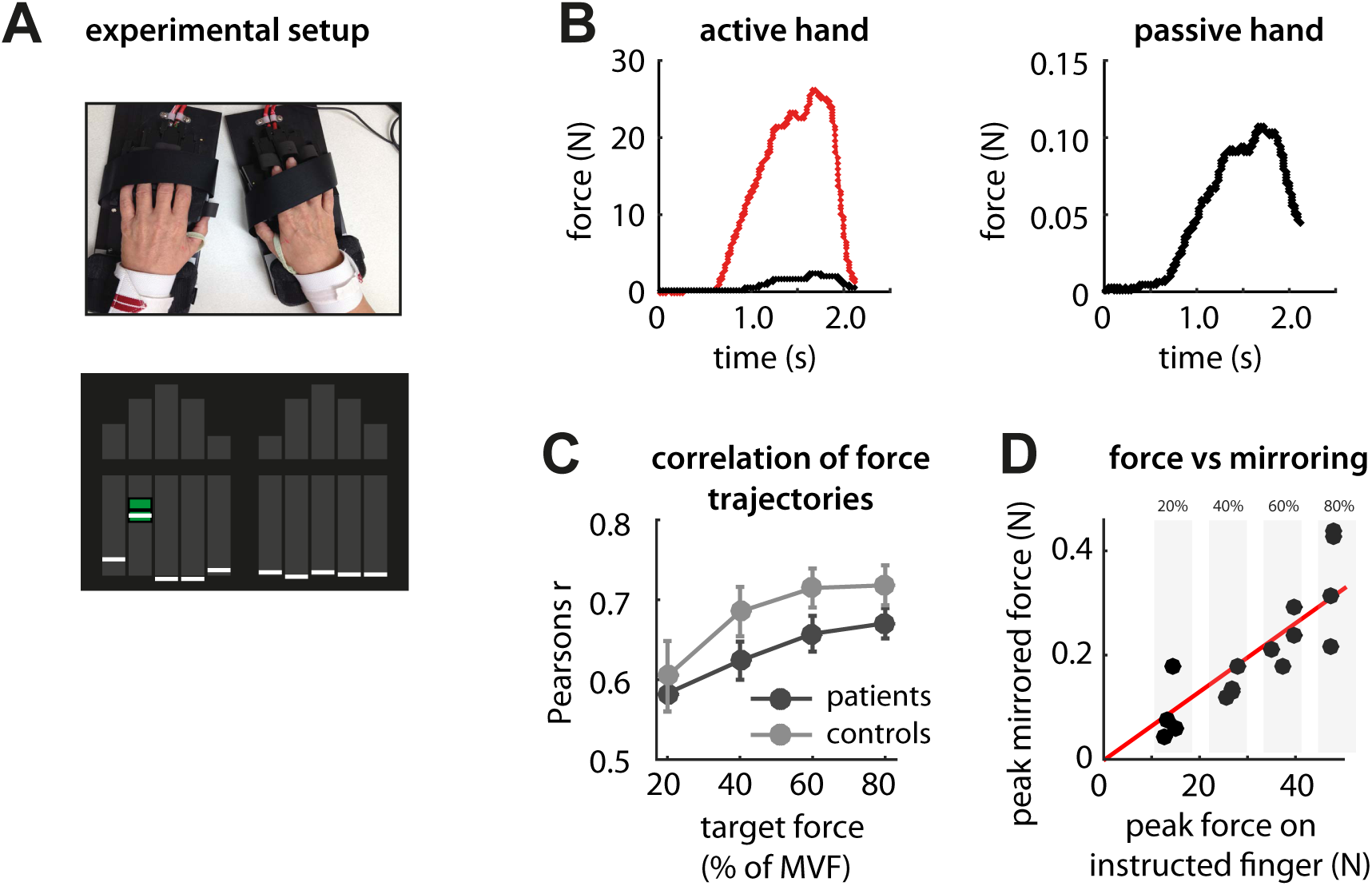
Assessment of mirror movements. (A) Both hands were strapped onto an ergonomic hand device capable of measuring isometric forces generated at the fingertips. Controls and patients were instructed to generate isometric forces by making individuated presses to bring the cursor into the target zone shown in green. During each measurement session, individuated finger presses were made at 20%, 40%, 60% and 80% of the maximum voluntary force on that finger. (B) Force presses with the instructed finger (thumb finger of right hand shown in red) resulted in involuntary forces on the passive fingers of the same hand (black), and subtle mirrored forces on the fingers of the passive hand (right panel). (C) Mirrored force trajectories were similar to that for the instructed finger, especially at higher target force levels. (D) Mirroring was quantified as the linear slope between the peak forces produced by the instructed finger and the peak averaged forces on the passive hand. The linear slope was log-transformed to allow the use of parametric statistical test, but for the purpose of clarity the raw values of the linear slope are reported in all subsequent figures.

14 neurologically-healthy participants were also recruited as healthy controls for the study (4 female; age=64.0, SD=8.2 years). Controls and patients did not differ in age (t_65_=1.60, p=0.11).

Data was collected across three centres: Johns Hopkins University, University of Zurich, and Columbia University. All experimental procedures were approved by the respective local ethics committee, and written consent was obtained from all participants.

### Apparatus to measure finger forces

We used a custom-built ergonomic keyboard (Fig. 1A) to measure isometric finger forces generated during the behavioural and fMRI tasks. During either experiment, participants were instructed to always keep both their hands on the 10 keys of the device. Force transducers beneath each key (Honeywell FS, dynamic range 0-25N) allowed for the sensitive measurement of finger forces in the instructed hand (Ejaz *et al*., 2015) (Fig. 1B), as well as mirrored finger forces in the passive hand (Diedrichsen *et al*., 2013).

### Assessment of mirror movements during the behavioural task

Mirror movements for each participant (patients and controls) were assessed over five longitudinal measurement sessions following recruitment (Table 1); weeks 2, 4, 12, 24 and 52 post-stroke.

**Table 1.** Patient information and measurement schedules for the behavioural and fMRI experiments. A total of 53 patients and 14 age-matched controls were recruited for the study and measured at five different time points over the course of a year. For the behavioural experiment, each participant in the study was on average measured over at least 3 sessions (patients, 3.5±1.5 sessions; controls, 4.3±1.4), with the overall experimental data being 70.1% complete for patients and 85.7% complete for controls. For the fMRI experiment, a subset of participants from the cohort were measured (N=12 controls and N=35 patients), with the experimental data being 73.7% complete for patients and 90% for controls.

| **week** | **2** | **4** | **12** | **24** | **52** |
| --- | --- | --- | --- | --- | --- |
| days (mean±SD) | 10 ± 4 | 37 ± 8 | 95 ± 10 | 187 ± 12 | 370 ± 9 |
| ***Behavioural experiment*** | **53 patients, 14 controls** |  |  |  |  |
| measured at week (%) |  |  |  |  |  |
| controls | 14 (100%) | 10 (71%) | 12 (86%) | 12 (86%) | 12 (86%) |
| patients | 39 (74%) | 39 (74%) | 40 (75%) | 39 (74%) | 31 (58%) |
| Fugl-Meyer (0.25-0.75 percentile) | (16-59) | (34-64) | (52-66) | (57-66) | (59-66) |
| ***fMRI experiment*** | **35 patients, 12 controls** |  |  |  |  |
| measured at week (%) |  |  |  |  |  |
| controls | 11 (92%) | 10 (83) | 11 (92%) | 11 (92%) | 11 (92%) |
| patients | 24 (69%) | 31 (89%) | 27 (77%) | 28 (80%) | 19 (54%) |
| Fugl-Meyer (0.25-0.75 percentile) | (16-60) | (45-65) | (59-65) | (60-66) | (64-66) |

During each measurement session, participants performed individuated force presses in the flexion direction with the instructed finger, while mirrored forces in the fingers of the passive hand were recorded. A visual representation of all ten fingers was presented on a screen (Fig. 1A). The experiment began by estimating the strength of each finger, measuring 2 repetitions of the maximum voluntary force (MVF) of each digit on both hands.

All subsequent trials required the production of isometric fingertip forces at a fraction of the MVF for the instructed digit (at 20%, 40%, 60%, 80%). At the start of every trial, a force target-zone (target-force±25%) on a single finger was highlighted in green. This was the cue for participants to make a short force press with the instructed finger to match and maintain the target-force for 0.5s. The trial was stopped if force on the instructed digit did not exceed 2.5N in the 2s following stimulus onset. Trials were presented in sequential order, starting from the left thumb to the left little finger, and ending with the right thumb to the right little finger. Trials were grouped as blocks, with each block consisting of one measurement each for the four target-force levels across the 10 fingers (4 target-force levels × 10 fingers=40 trials/block). Participant’s performed 4 such blocks during each measurement session.

### Quantifying the degree of mirror movements

During each trial, finger presses with the instructed finger resulted in subtle forces in the fingers of the passive hand (Fig. 1B). These mirrored forces were substantially smaller than the forces produced by the instructed finger. Even at the lowest target-force levels, the trajectory of these averaged mirrored forces correlated strongly with those produced by the instructed fingers (Fig. 1C). This was true for both controls (*r*=0.63, 95% confidence interval: 0.53-0.72), and patients (*r*=0.61, 95% confidence interval: 0.56-0.65). These correlations increased monotonically as the target-forces increased, consistent with previous reports that mirrored forces are a function of the force applied with the active hand (Armatas *et al*., 1996; Todor and Lazarus, 1986).

To quantify peak forces produced during mirroring, the resting baseline force on each finger prior to movement was subtracted from the subsequent force trace produced during the trial. Then the peak force *F*_*passive*_ on the passive hand was calculated as the peak averaged force on the fingers during the trial:

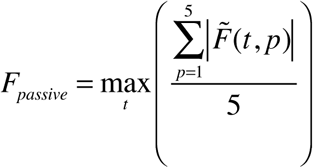

where *t* is the duration of the trial in seconds, and 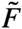 are the baseline corrected forces on finger *p* of the passive hand. Thus, *F*_*passive*_ indicates the peak averaged force in the passive hand when the active finger produces force.

The passive mirrored force increased approximately linearly with the force exerted by the active hand (Fig. 1D). To derive a singular metric of the degree of mirroring across the different target force levels, we conducted a regression analysis to estimate the ratio of the peak force on the instructed finger *F*_*active*_ and the peak mirrored force (*F*_*passive*_). First, all trials belonging to movements of the same instructed finger were grouped together. We plotted *F*_*active*_ on the x-axis and *F*_*passive*_ for corresponding trials on the y-axis and estimated the best-fit line forced through the origin that described the data points (Fig. 1D). Sensitivity to outliers was reduced by using robust regression with a b-squared weighting function. To ensure that the passive force was specific to mirroring and not due to spurious finger presses of the passive hand, we only used trials where the correlations between averaged force trajectories across all fingers in the active and passive hands were ≥0.2 to estimate the linear slope.

Finally, to allow for the use of parametric statistics, the regression slope (i.e. the estimate of the ratio) was log-transformed to make it conform better to a normal distribution. This log-slope provides a sensitive measure of mirroring in the passive hand due to movements of the instructed finger. For each participant, the log-slopes associated with the instructed fingers on each hand were averaged to get a composite metric of the degree of mirroring.

### Quantifying recruitment of fingers during mirror movements

The principle aim of this study was to determine how fingers if the passive hand were recruited during mirroring. To do so, we first calculated the mirroring across all 25 possible combinations of instructed/non-instructed finger pairs. Mirroring across each finger pair (*i*, *j*) was computed as described in the preceding section, by computing the log-slope between the peak force in the instructed finger *i*, and the peak force on the non-instructed finger *j*. The pattern of finger recruitment during mirroring was quantified separately for each participant and measurement session, thereinafter referred to as *mirroring pattern*.

To determine the degree of homologous mirroring, we averaged the log-slopes for homologous finger pairs (*i* = *j*) across the two hands for each participant. Non-homologous mirroring was determined by averaging log-slopes for all finger pairs where *i* ≠ *j*.

### Estimating changes in mirroring patterns over time

To estimate similarities between mirroring patterns for patients and controls, we first estimated the average mirroring pattern for all controls. This control pattern was then correlated with the corresponding mirroring pattern for each patient, separately for each week. The resulting correlations quantified the similarities between mirroring patterns for patients and controls during recovery. Since the mirroring patterns for controls were themselves estimated in the presence of measurement noise, even a perfect match between patient and control mirroring patterns would not result in a correlation of 1. To estimate a noise ceiling for the correlations, we calculated the average correlation of each controls mirroring pattern with the group mean. As a lower bound, each control’s mirroring pattern was also correlated with the group mean in which this participant was removed. These upper and lower bounds therefore specify the range of values correlations between mirroring patterns for control and patients could maximally take given measurement noise.

### Quantifying mirror movements in the paretic hand

In addition to the non-paretic hand, we also quantified the degree of homologous and non-homologous mirroring in the paretic hand during non-paretic finger presses. Since mirroring in the paretic hand might be influenced by the loss of hand strength, we restricted our analysis to a subset of relatively mildly impaired patients. Patients were split into a mild and severe group based on whether reliable muscle potentials could be evoked on the paretic hand during transcranial magnetic stimulation (TMS) of the lesioned hemisphere. Only TMS measurements obtained within the first 2 weeks after stroke were used to categorize patients. During each measurement session, 10 single TMS pulses were applied to the hand area of the motor cortex in the lesioned hemisphere while muscle activity from the contralateral FDI muscle was recorded. Patients that demonstrated reliable muscle evoked-potentials (MEP≥50*μ*V) for at least 5 out of the 10 TMS pulses were placed into the mild group, while those that did not show reliable MEPs even at 100% stimulation intensity were placed in the severe group. For the TMS experiment, only a subset of 40 patients (Fugl-Meyer, 16-59, 25%-75% percentile) were measured. Of these, 11 patients did not demonstrate reliable MEPs at week 2 and were thus categorized as severe, while 29 patients were categorized as mild and we focused our analysis on this subgroup.

### Quantifying finger individuation ability

In addition to the mirrored forces, individuated finger presses also resulted in enslaved forces on the uninstructed fingers of the active hand (Fig. 1B). These enslaved forces were generally much larger than the associated mirrored forces, and at high force requirements, degraded the participants ability to individuate a single finger (Z. M. Li *et al*., 1998). We quantified the degree of enslaving in the same way as for mirroring, by estimating the log-slope between the peak forces on the instructed and the passive fingers on the active hand respectively. We have previously used a similar metric to quantify patients impairment in finger individuation ability after stroke (Xu *et al*., 2016).

### Assessing neural activity associated with individuated finger movements (fMRI)

Cortical activity associated with finger movements was measured in controls and patients at the same time points as for the behavioural measurements, five times over the course of a 1-year period (Table 1).

Participants were instructed to produce individuated finger movements inside an MRI scanner in a protocol resembling the behavioural task. To reduce scanning time, only four fingers on either hand were tested (ring finger was excluded). Each trial required the production of 4 short isometric force presses with an instructed finger. Each trial began with the instructed finger highlighted in green for 2s. A green line then appeared below the finger stimulus as the go-cue for producing a short flexion force press with the instructed finger within 1.9s. This cue was repeated 4 times for a total of 4 repetitive presses with the instructed finger for that trial. A successful finger press required the production of either 1.8N or 8% of the MVF for that finger, whichever was lower. The green line turned blue to signal a successful finger press. Trials were grouped as experimental runs, with each run consisting of 3 trials for the 8 fingers across the two hands (a total of 3x8=24 trials/run). Trials within each run were presented in pseudo-random order, and participants performed 8 runs at each measurement session.

Functional scans during task performance were obtained at three centers on two different 3T Philips systems (Achieva and Ingenia). Scans were obtained with a 32-channel head-coil using a two-dimensional echo-planar imaging sequence (TR=2s, 35 slices, 154 volumes-per-run, slice thickness 2.5mm, 0mm gap, in-plane resolution 2.5x2.5mm^2^). Within each imaging run, six rest phases lasting 10s were randomly interspersed. A T1-weighted anatomical image (3D MPRAGE sequence, 1x1x1.2mm, 240x256x204mm FOV) was also acquired. For each participant, two diffusion tensor-imaging (DTI) images (TR=6.6s, 60 slices, 2.2mm slice thickness, 212x212mm FOV) were also acquired to help quantify the size and location of stroke lesions.

### Imaging analysis

All functional data was corrected for motion across runs (Diedrichsen and Shadmehr, 2005), and co-registered to the T1-image obtained in the participant’s first measurement session (either week 2 or 4). The raw time-series data was analyzed using a generalized-linear model (GLM) with a separate regressor for each finger/hand/imaging run (4-fingers × 2-hands × 8-runs = 64-regressors). Activation for each trial was modelled using a boxcar function (10.88s) convolved with a standard haemodynamic response function.

Each participants T1-image was used to reconstruct the pial and white-gray matter surfaces using Freesurfer (Dale *et al*., 1999). Individual surfaces were aligned across participants and registered to match a template using the sulcal-depth map and local curvature as minimization constraints.

The anatomical regions of interest (ROIs) were defined on the group surface using probabilistic cyto-architectonic maps aligned to the average surface (Fischl *et al*., 2008). Surface nodes with the highest probability for Brodmann area (BA4) 2cm above and below the hand-knob were selected as belonging to M1. Similarly, nodes in the hand-region in S1 were isolated using BA 3a, 3b, 1 and 2 (combined), again 2 cm above and below the hand knob.

Each participants DTI and T1-images (at first measurement) were used to estimate the size and location of lesions in two ROIs: i) cortical grey matter in the sensorimotor cortices (M1/S1) of either hemisphere, and the ii) corticospinal tract superior to the pyramids. Lesion boundaries were determined independently by radiologist (AVF) and neurologist (MB) that were blind to the patients clinical information and task performance. Detailed information about lesion distribution can be found in Xu et al. (2016).

Finally, the parameter estimates from the GLM analysis in M1 and S1 ROIs with lesion areas excluded, were identified and pre-whitened using the GLM residuals to reduce the effects of estimation noise (Walther *et al*., 2015). These pre-whitened parameter estimates quantified the evoked-BOLD activations. As measuring participant data for all 5 sessions was ambitious, we ended up with an unbalanced experimental design due to missing data across the fMRI experiment. We therefore used linear mixed-effects models for the summary plots of the fMRI experiment (Fig 5D; *lme4* package in R; (Bates *et al*., 2014)) to account for the problem of missing values.

### Statistical analysis

We used 2-sided t-tests to test for differences in means either across groups, or across different time-points of recovery. To test for differences between summary statistics across groups or over time, we used linear mixed-effects models in the lme4 package in R (Bates *et al*., 2014). In all statistical models, an intercept was included as one of the fixed effects, with each participant considered a random-effect. All data presented in the text and figures are represented as mean±standard error of the mean. All statistical tests involving correlations were performed on Fisher Z-transformed values.

## Results

### Mirror movements appeared early after stroke and normalized over the year

Using a sensitive behavioural assay, we quantified mirror movements in 53 stroke patients and 14 controls. The first measurement was within the first 2-weeks of stroke-onset, and subsequently at four sessions over the following year (Table 1). During each measurement session, patients and controls produced individuated finger presses at different target-force levels while forces in the passive hand were measured (Fig 2B). To quantify the degree of mirroring, we calculated the linear slope between the peak force produced by the instructed finger and the peak averaged force in the passive hand (Fig 1D; see methods).

**Figure 2.**
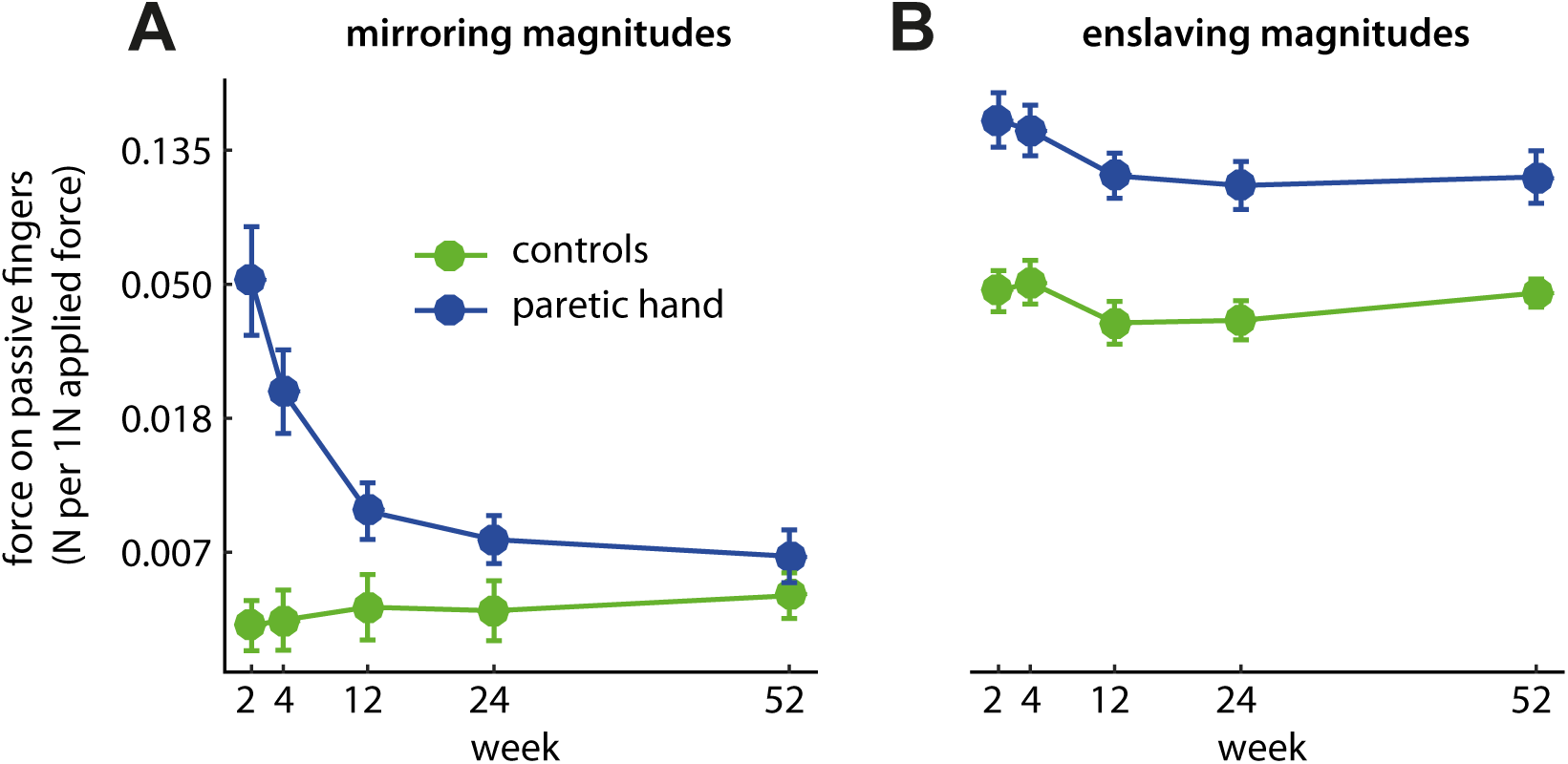
Longitudinal changes in mirror movements and fine-finger control after stroke. (A) Changes in mirroring for controls and patients measured over the course of a year. For patients, mirroring was measured in the fingers of the non-paretic hand, during active finger presses with the paretic hand. (B) Associated changes in fine-finger control on the active hand across groups. Individuated finger presses in patients and controls resulted in undesired force contractions on the uninstructed fingers of the active hand. The larger these so-called enslaved movements, the worst the degree of fine-finger control. For clarity, the raw values of the linear-slope estimates for mirroring and enslaving are plotted in (A) and (B).

Patients showed large time-course changes in mirroring in the year following a stroke (Fig. 2A). In the first two weeks after damage (week 2), individuated finger presses with the paretic hand resulted in large forces in the non-paretic hand, with 1N of voluntary force resulting in approximately 0.051N of averaged mirrored force. In comparison, mirroring in controls was significantly lower than patients (1N/0.004N; *t*_51_=3.67, *p*=0.001). Mirroring in patients subsequently reduced over time (*χ*^2^=82.99, *p*<<0.0001). However, even 6-months after stroke, mirroring was still marginally larger in comparison to controls (*t*_51_=1.75, *p*=0.087). There was a strong correlation between mirroring during the early and late stages following stroke *r*=0.73 (*p*<0.001), demonstrating that patients who exhibited large mirroring early after stroke continued to do so throughout recovery.

The longitudinal changes in mirroring were remarkably similar to those for the deficits in fine-finger function in the paretic hand (Fig. 2B). After stroke, patients’ efforts to produce isometric forces with a single finger resulted in abnormally large forces in the uninstructed fingers of the paretic hand. These enslaved forces signify the loss of fine-finger control in patients (S. Li *et al*., 2003; Xu *et al*., 2016). Early after damage (week 2), enslaving in patients was significantly larger than controls, demonstrating a substantial loss of individuated finger control (controls 0.042N/1N; patients 0.170N/1N; t_51_=4.02, p<0.001). Enslaving progressively reduced over the course of the year (χ^2^=28.38, p<<0.0001), but never fully normalized even by 6 months post stroke (t_51_=3.09, p=0.003). Patients who had large enslaving early after stroke also demonstrated large mirroring at the same time-period (enslaving and mirroring at week 2, r=0.78, p<<0.0001), and continued to do so even by the chronic stage of recovery (enslaving week 2 and mirroring week≥24, r=0.66, p=0.0001).

Consistent with earlier findings, here we found that mirroring in the non-paretic hand was exaggerated after stroke (Y. H. Kim *et al*., 2003; Nelles *et al*., 1998; Wittenberg *et al*., 2000), and appeared with a time-course that mimicked that for the fine-control deficits in the paretic hand.

### Mirror movements were characterized by the recruitment of multiple fingers

Next, we were interested in understanding finger recruitment patterns in the non-paretic hand during mirror movements. Specifically, we wanted to determine the extent to which mirroring in the non-paretic hand was characterized by a broad recruitment of fingers. We therefore characterized mirroring patterns across all active/passive fingers in both controls and patients (see methods).

The degree of mirroring in each passive finger as a function of the instructed finger can be seen in Figure 3A. The overall patterns of mirroring across all active/passive finger pairs themselves were highly reliable, with split-half correlations being r>0.85 for both controls and patients (Supplementary Table 1). The first immediate observation is that mirroring was not restricted to the homologous fingers (diagonal), but that substantial effects could also be observed on non-homologous fingers (off-diagonal). To quantify this observation, we partitioned mirroring across the different active/passive finger pairs into their respective homologous and non-homologous components (see methods).

**Figure 3.**
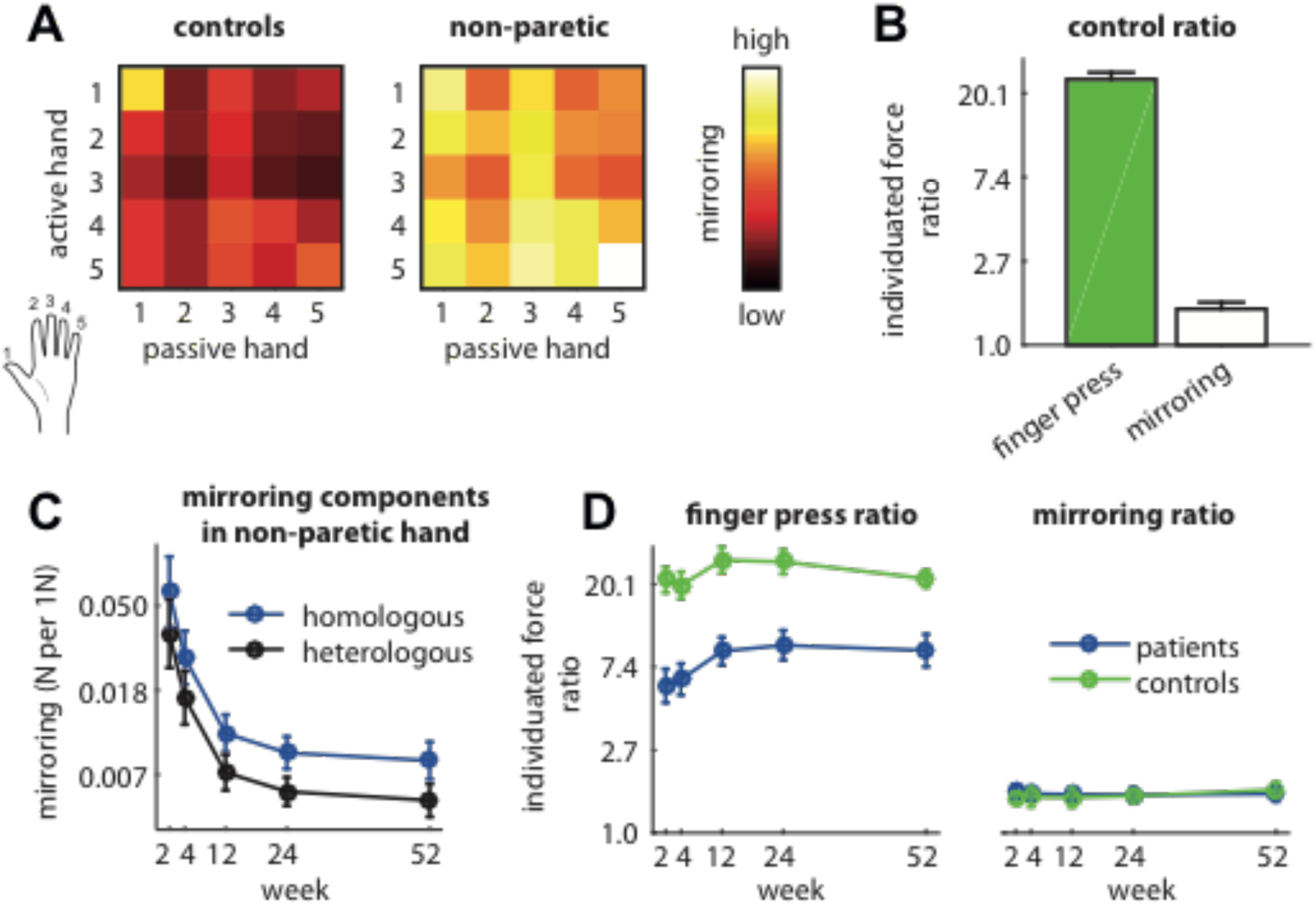
Relative contributions of homologous and non-homologous components to mirror movements on the non-paretic hand. (A) Mirroring across all possible active/passive finger pairs for controls and patients (on non-paretic hand only). Rows and columns denote which finger was pressed on the active hand, and the finger on the passive hand that mirroring was estimated on, respectively. Diagonal and off-diagonal matrix entries represent mirroring across homologous and non-homologous finger pairs. (B) Individuated finger presses by controls resulted in enslaved forces on the passive fingers of the same hand and mirrored forces across homologous and non-homologous finger pairs. The ratio between instructed/enslaved forces within the active hand is shown in green, while ratio between homologous and non-homologous mirroring components is shown in white. (C) Changes in homologous and non-homologous mirroring components on the non-paretic hand in the year following stroke. For clarity, the raw values of the linear-slope estimates for mirroring are plotted. (D) For patients, the ratios between instructed/enslaved forces on the paretic hand, and the ratio between homologous/non-homologous mirroring patterns are shown in the left and right panels respectively.

In controls, finger presses resulted in a broad recruitment of fingers in the passive hand. Finger presses in the active hand were highly individuated in nature, with 1N of force on the instructed finger resulting in 0.042N of enslaved forces (ratio of 24.77±2.18; Fig. 2B). These finger presses resulted in mirroring across both homologous and non-homologous fingers pairs. While homologous mirroring was, on average, larger than the non-homologous component (t_13_=5.421, p=0.0001), some finger presses resulted in near equivalent effects on both (index finger presses; t_13_=1.23, p=0.240, ring; t_13_=0.88, p=0.398). Overall, forces in the passive hand were much more evenly distributed across fingers than the forces in the active hand (Fig. 3B), with the corresponding ratio between homologous and non-homologous mirroring components (1.61±0.16) being nearly 15 times smaller than the instructed/enslaving ratio on the active hand (t_13_=28.26, p<<0.0001). Thus, mirroring was not simply due to a symmetric digit-by-digit activation of the motor system, as predicted from the exact mirroring of cortical activity patterns across hemispheres (Diedrichsen *et al*., 2013; Liu *et al*., 2010; Scherer *et al*., 2009).

Similarly, in patients, finger presses with the paretic hand resulted in a broad recruitment of fingers in the non-paretic hand. The year-long changes in mirroring characterized earlier (Fig. 2A) were observed in both homologous and non-homologous fingers (Fig. 3C; change over weeks: homologous, χ^2^=71.35, p<<0.0001, non-homologous, χ^2^=78.15, p<<0.0001), with homologous mirroring being the stronger of the two (χ^2^=24.53, p<<0.0001). Critically, despite these longitudinal changes, the ratio between homologous and non-homologous mirroring (1.76±0.12) remained stable across weeks (χ^2^=1.16, p=0.885) and was at the same level as healthy controls (χ^2^=0.10, p=0.754).

To summarize, finger presses in patients, like controls, resulted in a broad recruitment of fingers in the passive hand. Remarkably, when considering mirroring across all active/passive fingers irrespective of the homologous and non-homologous finger (Supplementary Figure 2), a high degree of similarity between finger recruitment patterns for patients and controls was observed. Throughout recovery, mirroring patterns for patients looked like a scaled version of the corresponding control mirroring pattern. The most parsimonious explanation for this similarity would be that a single system is responsible for mirroring in controls, and it is (un)up-regulated in the non-paretic hand after stroke. However, in the next section, we characterize mirror movements in the paretic hand and provide evidence that more than one system appears to contribute towards mirroring.

### Homologous and non-homologous mirroring dissociated in the paretic hand

After stroke, not only is mirroring exaggerated in the non-paretic hand, but a slight reduction of mirroring in the paretic hand is also observed during non-paretic hand movements (Nelles *et al*., 1998). Mirroring in the paretic hand has to-date received little attention, consequently the cause for reduced mirroring is unknown. We hypothesized that if homologous mirroring is primarily contributed by cortical motor areas, then stroke-related damage in the lesioned hemisphere should result in reduced mirroring in the primarily the homologous fingers of the paretic hand. To test this, we partitioned mirroring across all active/passive finger pairs into their respective homologous and non-homologous components.

Since the degree of mirroring in the paretic hand can be influenced by a loss of hand strength, we restricted our analysis to a subgroup of mild patients who demonstrated reliable muscle-evoked potentials early after stroke (see methods). Even in the early period after stroke, these mild patients had sufficient residual strength to express mirroring at the level of controls (Fig. 4A; 1N/0.004N). Infact, even at maximal force production with the non-paretic hand (15.7N), the predicted mirrored forces on the paretic hand were small (0.07N) in comparison to the residual strength on the hand (9.0N; residual strength versus predicted mirroring at control level, t_21_=6.77, p<<0.0001). Thus, these mild patients had sufficient strength to exhibit mirroring in the paretic hand.

**Figure 4.**
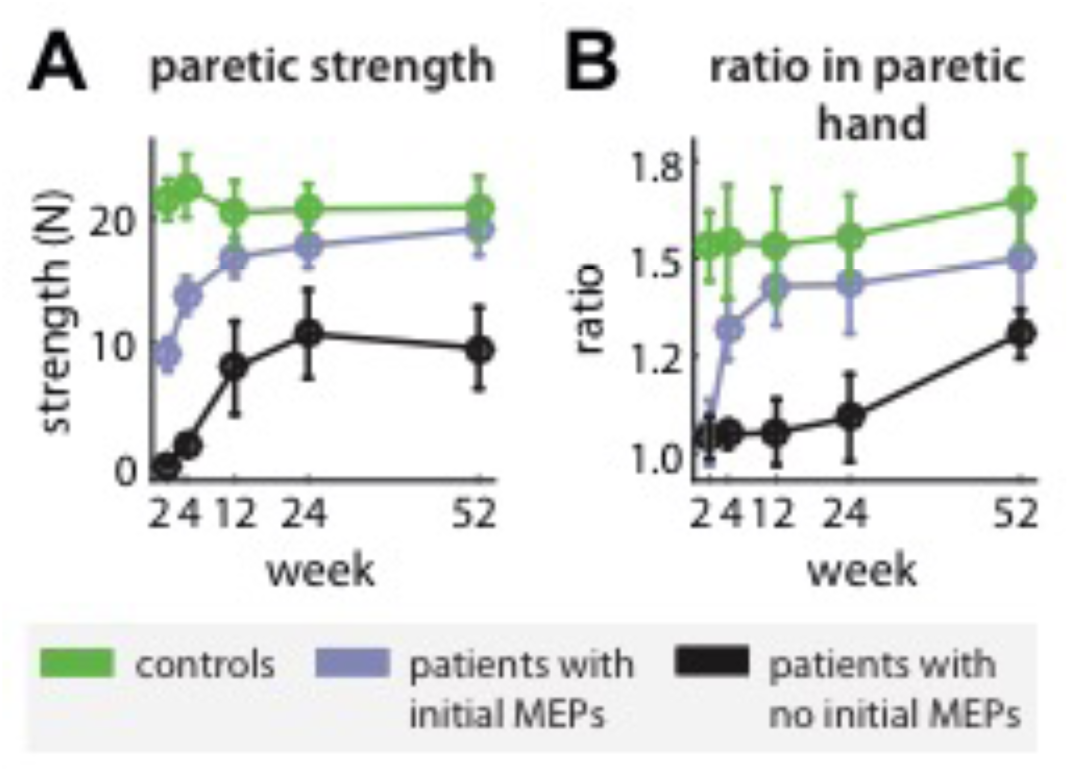
The homologous and non-homologous components of mirror movements in the paretic hand. (A) Time course of strength recovery in patients who demonstrated reliable MEPs (mild group) in the first few weeks after stroke, and those who did not (severe group). (B) Ratios between the homologous and non-homologous mirroring components across the mild and severe groups.

However, as predicted, the ratios between homologous and non-homologous mirroring was approximately equal early after stroke (Fig. 4B; week 2; ratio for mild group=1.11±0.11). Mirroring subsequently became stronger in the homologous finger pairs as the paretic hand regained fine-finger function, with the homologous/non-homologous ratio progressively increasing during recovery (χ^2^=21.47, p=0.0003), eventually normalizing to the control level (week≥24; t_36_=0.48, p=0.632). This reduction in the homologous component of mirroring was also seen for the severe patients (weeks≤24; Supplementary Figure 3), with the ratio between homologous and non-homologous remaining low even though strength recovered during this time.

To summarize, we demonstrate that the homologous and non-homologous components of mirroring in the paretic hand appear to dissociate, despite patients having sufficient strength in the hand. This dissociation effect is hard to attribute to a single system contributing towards mirroring. We therefore conclude that mirror movements after stroke are generated by contributions from (at least) two separate systems.

### No modulation of evoked-BOLD activities in the bilateral sensorimotor cortices after stroke

Finally, we consider the neurophysiological mechanisms that could cause an exaggeration of mirror movements in the non-paretic hand after stroke. One candidate mechanism could be the large activations previously reported in the primary somatosensory (S1) and motor (M1) cortices of the non-lesioned hemisphere after stroke (Cincotta and Ziemann, 2008; Cramer *et al*., 1997; Y. H. Kim *et al*., 2003; Ward *et al*., 2003; Wittenberg *et al*., 2000). These activations could potentially exaggerate mirroring directly or indirectly. In the first case, activations could be directly transmitted to the motoneurons/spinal interneurons that control the passive hand, via the crossed corticospinal pathway. Alternatively, the activations could indirectly exaggerate mirroring by up-regulating the activity of subcortical motor circuits through cortico-brainstem connections (Fisher *et al*., 2012).

If mirror movements after stroke were caused by over-activation of the non-lesioned sensorimotor cortex, then the time-course of these activations should resemble the time-course changes in mirroring quantified earlier (Fig. 2A). To test this idea, we used fMRI to measure evoked-activities in the hand area of S1/M1, in a smaller subset of participants from the same study cohort (Table 1, 35 patients, 12 controls). Participants performed individuated finger presses inside an MRI scanner (Fig. 5A). During paretic finger presses, patients demonstrate the same mirroring and enslaving behaviour both inside and outside the scanner environments (Fig. 5B-C; mirroring, r=0.89, p<<0.001; enslaving, r=0.75, p<<0.001).

**Figure 5.**
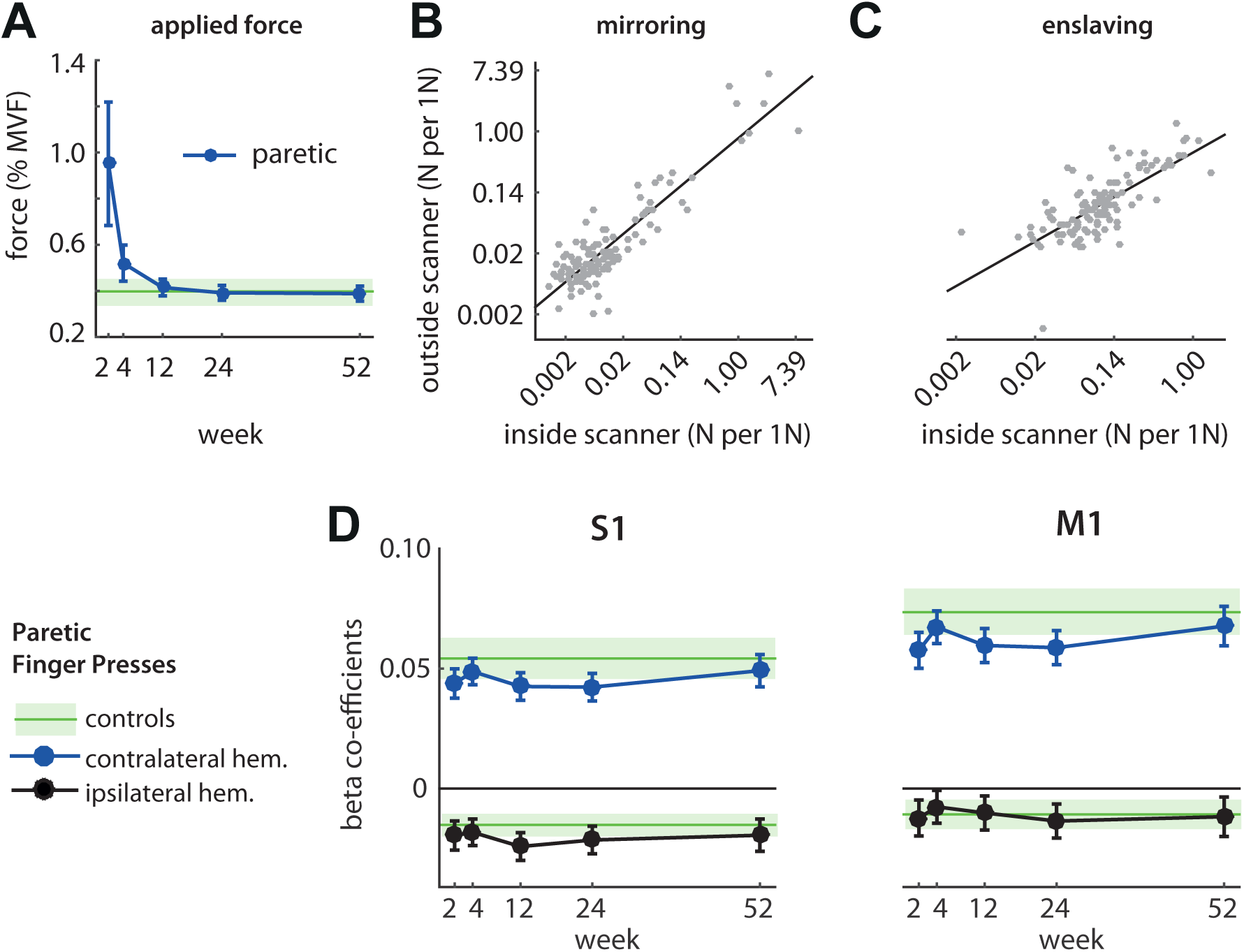
Evoked-BOLD activities for finger presses in the primary somatosensory (S1) and motor (M1) cortices. (A) During the fMRI task, patients and controls were required to produce either 1.8N or 8% of the maximum voluntary force (MVF) on the finger. Forces are expressed as a percentage of MVF. Controls produced forces at approximately 40% of MVF. From week 4 onwards, forces produced by patients and controls were not significantly different (week≥4; χ^2^=0.02, p=0.887). (B) Measurements of mirroring on the non-paretic hand were highly correlated inside and outside the scanner environments. (C) Similarly, enslaving in the paretic hand was highly correlated for measurements inside and outside the scanner environments. Each dot in B-C represents the session measurement of a single patient. For clarity, the raw values of the linear-slope estimates for mirroring are plotted in (B-C). (D) Evoked-BOLD activities in contra- and ipsilateral S1 and M1 cortices due to paretic finger presses. Corresponding contra and ipsi activities in controls are depicted by the shaded green regions (Mean±SE).

The resulting evoked BOLD responses in M1/S1 for patients were remarkably stable throughout recovery (Fig. 6D; statistics in Table 2). For paretic hand presses, we did not find any time-course related changes in the evoked-activities in either the contra- or the ipsi-lateral cortices, with activations in either hemisphere indistinguishable from their counterpart in controls. Patients continued to demonstrate the stereotypical pattern of evoked cortical responses seen for unimanual finger presses in health, which was characterized by an increase and reduction of BOLD responses in the contra- and ipsilateral sensorimotor cortices respectively.

**Table 2.** Statistics for the fMRI experiment. Statistics are shown for differences in contralateral and ipsilateral M1/S1 activations, across weeks (first two columns) and between patients and controls (last two columns).

|  | change over weeks |  | patients versus controls |  |
| --- | --- | --- | --- | --- |
| | $\chi^2$ | p | $\chi^2$ | p |
| activity for paretic presses |  |  |  |  |
| contra (S1) | 1.410 | 0.842 | 1.160 | 0.282 |
| contra (M1) | 2.070 | 0.723 | 1.150 | 0.285 |
| ipsi (S1) | 1.860 | 0.761 | 0.813 | 0.367 |
| ipsi (M1) | 1.250 | 0.870 | 0.010 | 0.915 |

To summarize, we report that the clear occurrence of the longitudinal changes in mirroring after stroke were not accompanied by over-activations in the sensorimotor cortices of either the non-lesioned or the lesioned hemispheres.

## Discussion

In this study, we present a detailed characterization of mirror movements that appear after stroke. Consistent with earlier findings, mirroring was exaggerated in the non-paretic hand (Y. H. Kim *et al*., 2003; Nelles *et al*., 1998; Sehm *et al*., 2009; Wittenberg *et al*., 2000). We expand upon these previous studies and demonstrate that mirroring appeared early after stroke and normalized as the hand recovered function. Despite these time-course changes in mirroring, we did not find any over-activations in the sensorimotor cortices in either hemisphere. These sensorimotor areas (M1/S1) provide the bulk of the inputs to the corticospinal pathways that provide fine-finger control (Lemon, 2008; Porter and Lemon, 1993), and the lack of evoked-BOLD modulation in these areas suggests that a simple up/down regulation of overall activity is unlikely to be the mechanism which exaggerates mirroring after stroke. Although, we cannot completely rule out the possibility that BOLD responses might be insensitive to subtle changes in sensorimotor activity required to produce the small forces during mirroring, our results contradict earlier studies that have argued that exaggerated non-paretic mirroring is caused by over-activations in ipsi- or contralesional M1/S1 (Cincotta and Ziemann, 2008; Y. H. Kim *et al*., 2003; Wittenberg *et al*., 2000).

The main goal of this study was to better understand finger recruitment patterns during mirror movements after stroke. We did this by quantifying the distribution of mirrored forces across homologous and/or non-homologous fingers, attributing homologous finger forces to cortical pathways, while attributing broad distribution of forces across all fingers to subcortical pathways instead. Our approach is analogous to the recent approach by Dean and Baker (2016) who investigated reticular contributions towards hand function using the StartReact paradigm (Valls-Solé *et al*., 1995). The authors compared muscle activations in the hand during the presentation of intense and mild acoustic cues, predicting that intense acoustic cues would preferentially rely on contributions from the reticulospinal system and therefore elicit less fractionated muscle activity when compared to milder cues. While the authors reported no differential effect of startling acoustic cues on hand muscle activity, here we report that the distribution of mirrored forces on the passive hand are indeed less fractionated than would be predicted by the forces on the active hand that generated them. By quantifying finger recruitment patterns during mirroring in both the non-paretic and the paretic hand, we find evidence of two components of mirroring, with the two components characterized by a broad recruitment of fingers, and recruitment of the homologous finger respectively. The first mirroring component (broad finger recruitment) has to-date remained undocument, primarily because previous studies have only focused on the homologous muscles/fingers (Armatas *et al*., 1994; Y. Kim *et al*., 2015; Koerte *et al*., 2010; Mayston *et al*., 1999). Our results therefore add to our current understanding of mirroring, both in stroke and health.

If the neocortex provides the ability to perform fine-finger function, then what should cortical contributions to mirror movements look like? Using data from recent fMRI studies, we argue that cortical activation patterns evoked during individuated finger presses predict mirroring primarily in the homologous finger of the passive hand. Specifically, individuated finger presses result in evoked-activities from motor areas distributed across the cortex (e.g. M1/S1, but also supplementary and premotor areas) (Diedrichsen *et al*., 2013; Ejaz *et al*., 2015). However, the activation patterns for a finger press are highly similar across the various cortical motor areas (e.g. M1; Pearson’s r=0.8) *regardless of whether the contralateral, or the homologous finger in the ipsilateral hand was used* (Diedrichsen *et al*., 2013). To the extent that these activation patterns specify the pattern of recruitment of muscles/fingers of the hand (Ejaz *et al*., 2015), cortical contributions to mirroring should primarily recruit the homologous passive finger.

Although subcortical contributions towards hand function in primates has been investigated in detail (Baker, 2011b; Lawrence and Kuypers, 1968b; Riddle *et al*., 2009; Soteropoulos *et al*., 2012; Zaaimi *et al*., 2012), the extent to which these subcortical pathways contribute towards hand function, and indeed mirror movements, in humans remains to be determined. One clue comes from comparing the patterns of upper-limb muscle recruitment during mirroring in humans, with muscle responses measured following stimulation of subcortical pathways in primates. For instance, in young children, flexion of the elbow joint results in mirroring mostly on the extensor muscles of the opposing elbow (Missiuro, 1963). This recruitment of ipsilateral flexors and contralateral extensor shoulder muscles is a prominent muscle activity pattern observed during stimulation of neurons in the ponto-medullary reticular formation (Herbert *et al*., 2010; Hirschauer and Buford, 2015). These neurons provide input to the reticulospinal tract which has been strongly implicated as a parallel pathway involved in hand function (Baker, 2011a; Riddle *et al*., 2009; Soteropoulos *et al*., 2012) and can therefore serve as a subcortical pathway capable of contributing towards mirroring.

If recovery of paretic hand function relies increasingly on the capacity of the subcortical systems to compensate for cortical damage (Xu *et al*., 2016), and if these pathways are responsible for contributing towards mirror movements, then how does mirroring reduce over the same time while paretic hand function recovers? Recent evidence from a primate study suggests that an increased reliance on bilaterally organized subcortical systems for paretic hand recovery, can in fact occur alongside a concomitant decrease in mirroring in the non-paretic hand. In the study, neurons in the ipsi- and contralateral sections of the ponto-medullary reticular formation (PMRF) were shown to alter the strength of their outputs onto motoneurons/spinal interneurons in either half of the spinal cord independently (Herbert *et al*., 2015). Specifically, connections between the paretic hand and cells in ipsi-PMRF were strengthened, while connections between the non-paretic hand and ipsi-PMRF cells were weakened. Therefore, such a pattern of subcortical reorganization could simultaneous facilitate recovery of the paretic hand and reduce the degree of mirroring in the non-paretic hand.

In conclusion, we have provided a detailed characterization of both the time-course and pattern of mirror movements following stroke. While mirroring is itself an interesting phenomenon that appears after damage, we propose that it additionally offers a window into cortical and subcortical contributions towards hand recovery.

## Acknowledgements

We would like to thank the tireless work of the many therapists and research associates that helped in the different facets of this project. We would also like to thank the patients for their valuable time and effort.

## Supplementary Material

**Supplementary Table 1.** Split-half reliabilities for mirroring patterns estimated across weeks. To estimate the reliability, data from each measurement session was dividing into odd and even runs, and the corresponding mirroring patterns for each half were estimated independently. Pearson’s correlation between the patterns from the two halves was then calculated to obtain the within-session or split-half reliability.

| week | 2 | 4 | 12 | 24 | 52 |
| --- | --- | --- | --- | --- | --- |
| ***controls*** |  |  |  |  |  |
| mean | 0.88 | 0.89 | 0.90 | 0.89 | 0.92 |
| range | (0.82-0.92) | (0.85-0.92) | (0.85-0.94) | (0.84-0.93) | (0.88-0.94) |
| ***non-paretic hand*** |  |  |  |  |  |
| non-paretic hand | 0.86 | 0.89 | 0.89 | 0.90 | 0.89 |
| range | (0.82-0.89) | (0.86-0.92) | (0.86-0.91) | (0.86-0.92) | (0.85-0.92) |
| ***paretic hand*** |  |  |  |  |  |
| paretic hand | 0.88 | 0.87 | 0.87 | 0.90 | 0.87 |
| range | (0.83-0.90) | (0.84-0.90) | (0.84-0.90) | (0.88-0.92) | (0.82-0.91) |

**Supplementary Figure 1.**
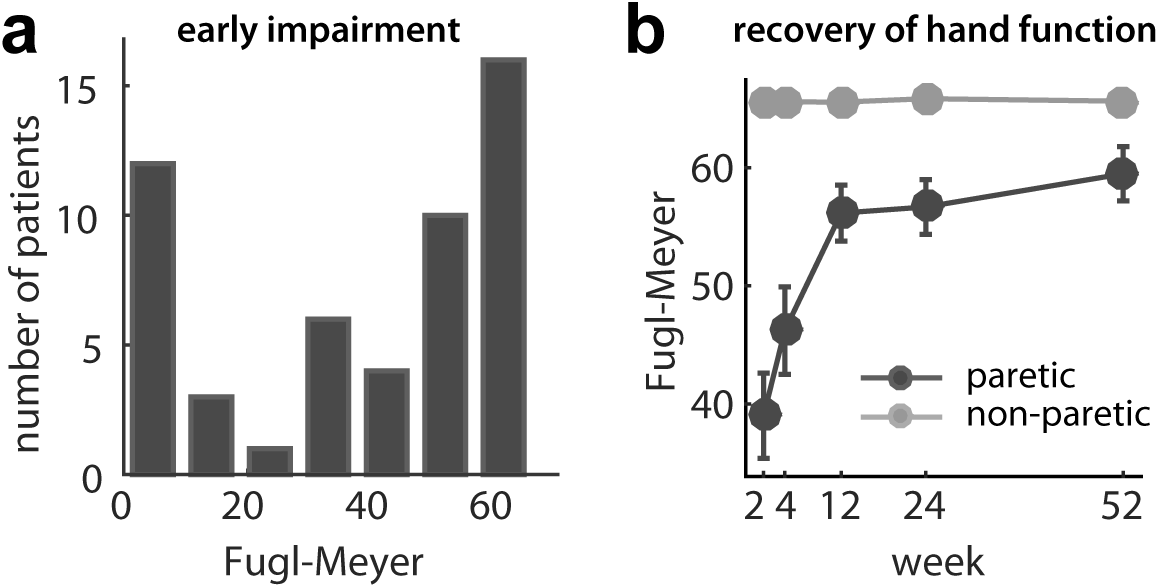
Patient information. (a) Distribution of Fugl-Meyer measurements on paretic hand at the point of first measurement (either week 2 or 4). (b) Fugl-Meyer measurements for patients over the course of 1 year following stroke.

**Supplementary Figure 2.**
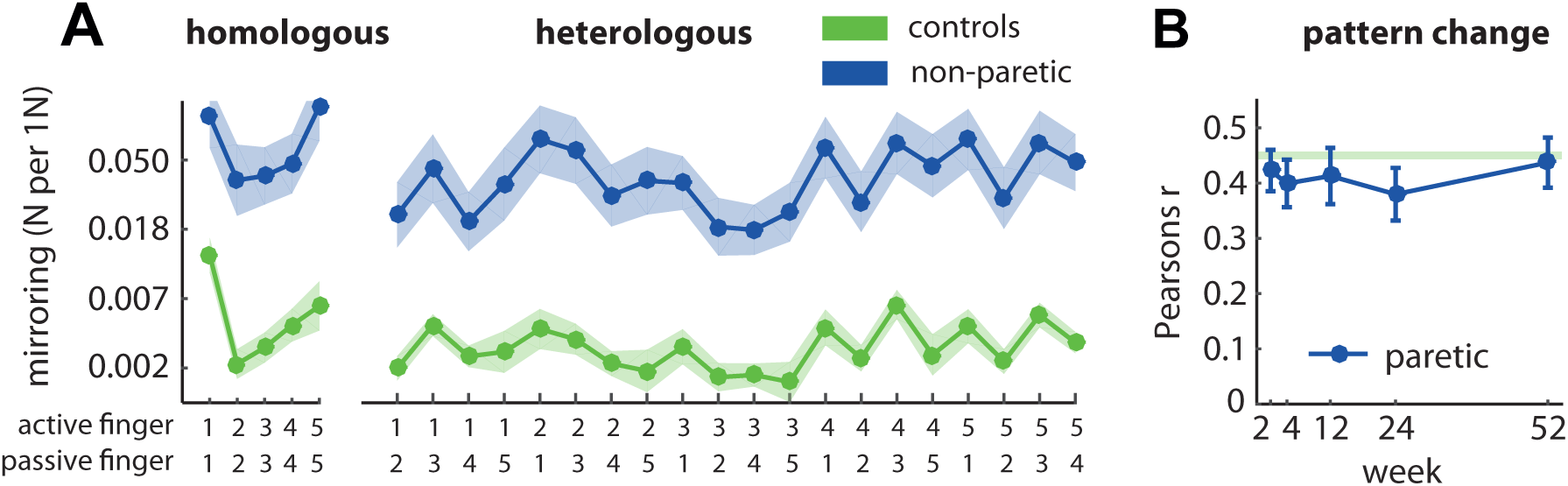
Stability of mirroring pattern during stroke recovery. (A) The average mirroring patterns across all active/passive finger pairs are shown for patients (week 2) and controls. For clarity, the raw values of the linear-slope estimates for mirroring are plotted in A. Similarity between the patterns for patients and controls was high, even in the early period after stroke (week 2, r=0.88, p<<0.0001). (B) Correlations between mirroring patterns for patients and controls remained unchanged throughout recovery (χ^2^=1.87, p=0.760). The pattern correlations for patients and controls were also close to noise ceilings; i.e. the maximum possible pattern correlations possible given the measurement noise on mirroring patterns for each control (see Methods).

**Supplementary Figure 3.**
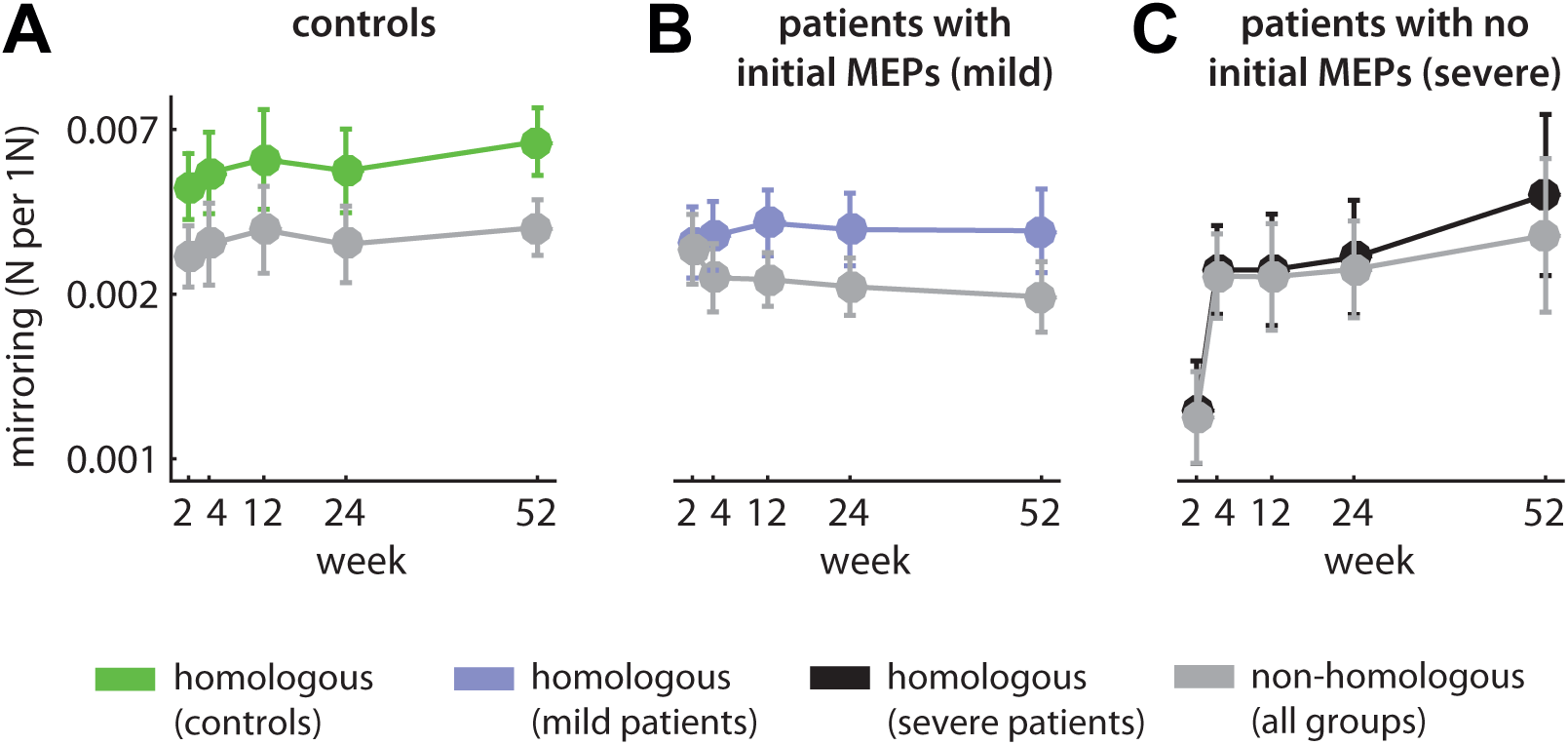
The homologous and non-homologous components of mirror movements during recovery in the (A) control group, (B) in patients who demonstrated reliable MEPs at weeks≤4 (mild group), and (C) in patients who not demonstrate reliable MEPs at weeks≤4 (severe group).

